# The role of *Caulobacter* cell surface structures in colonization of the air-liquid interface

**DOI:** 10.1101/524058

**Authors:** Aretha Fiebig

## Abstract

In aquatic environments, *Caulobacter* spp. are often present at the boundary between liquid and air known as the neuston. I report an approach to study temporal features of *Caulobacter crescentus* colonization and pellicle biofilm development at the air-liquid interface, and have defined the role of cell surface structures in this process. At this interface, *C. crescentus* initially forms a monolayer of cells bearing a surface adhesin known as the holdfast. When excised from the liquid surface, this monolayer strongly adheres to glass. The monolayer subsequently develops into a three-dimensional structure that is highly enriched in clusters of stalked cells known as rosettes. As this pellicle film matures, it becomes more cohesive and less adherent to a glass surface. A mutant strain lacking a flagellum does not efficiently reach the surface, and strains lacking type IV pili exhibit defects in organization of the three-dimensional pellicle. Strains unable to synthesize holdfast fail to accumulate at the boundary between air and liquid and do not form a pellicle. Phase contrast images support a model whereby the holdfast functions to trap *C. crescentus* cells at the air-liquid boundary. Unlike the holdfast, neither the flagellum nor type IV pili are required for *C. crescentus* to partition to the air-liquid interface. While it is well established that the holdfast enables adherence to solid surfaces, this study provides evidence that the holdfast has physicochemical properties required for partitioning of non-motile mother cells to the air-liquid interface, which facilitates colonization of this microenvironment.

**Importance:** In aquatic environments the boundary at the air interface is often highly enriched with nutrients and oxygen. Colonization of this niche likely confers a significant fitness advantage in many cases. This study provides evidence that the cell surface adhesin known as a holdfast enables *Caulobacter crescentus* to partition to and colonize the air-liquid interface. Additional surface structures including the flagellum and type IV pili are important determinants of colonization and biofilm formation at this boundary. Considering that holdfast-like adhesins are broadly conserved in *Caulobacter* spp. and other members of the diverse class *Alphaproteobacteria*, these surface structures may function broadly to facilitate colonization of air-liquid boundaries in a range of ecological contexts including freshwater, marine, and soil ecosystems.

## Introduction

In aqueous systems, macronutrients partition to and accumulate at surfaces at both solid-liquid and air-liquid boundaries (1, 2), and dissolved oxygen levels are highest at air interfaces. An ability to take advantage of elevated concentrations of nutrients and/or oxygen at such surface boundaries likely confers a significant growth advantage in many cases (3). Certainly, bacteria have long been noted to partition to submerged solid-surfaces (4, 5) and to air-liquid interfaces (6). Diverse morphological and metabolic characteristics of bacterial cells enable colonization of surface microenvironments.

As aquatic systems cover the majority of our planet, microbial activity in surface films has a significant impact on global biogeochemical cycles (7-10). Moreover, ecologically important aqueous interfaces are also found in terrestrial soils, where microbes primarily occupy the aqueous phase at solid- and air-liquid boundaries (10, 11). In porous soils and highly aerated bodies of water, bubbles provide mobile air-liquid surfaces upon which bacteria can be transported (10, 11). Though biofilms at air-liquid interfaces are not as well studied as solid surfaces, common themes in biofilm development in many species on varied surfaces have emerged over the past two decades. For example, flagellar motility and extracellular polysaccharides are important for colonization of both solid surfaces and air-liquid interfaces. In many cases, protein polymers such as pili and curli, or extracellular DNA also play a role in surface attachment and/or biofilm development (for reviews see (12-17)).

### A dimorphic bacterial model system to study colonization of the air-liquid interface

*Caulobacter* spp. are found in nearly any environment that experiences extended periods of moisture including marine, freshwater, and soil ecosystems (18, 19). Poindexter previously reported an approach to enrich *Caulobacter* spp. by sampling from the air-liquid interface (20). Specifically, she noted that when natural water samples are left to stand, a pellicle enriched with prosthecate (i.e. stalked) bacteria will form at the surface where liquid meets the air. *Caulobacter* have a dimorphic life cycle characteristic of many *Alphaproteobacteria* whereby each cell division yields a motile newborn swarmer cell and a sessile mother cell (20-22). In the case of *Caulobacter*, the sessile mother cell has a polar prosthecum, or stalk, while the swarmer cell has a single flagellum and multiple type IV pili at one cell pole. The swarmer cell further has the capacity to secrete a polar adhesin, called a holdfast, at its flagellated/piliated pole (23-25). Cells are motile for only a fraction of the cell cycle; swarmers transition to sessile stalked cells upon initiation of DNA replication and thus undergo a transition from motile to sessile with every round of cell division.

As a swarmer cell transitions to a stalked cell, the flagellum is shed, and the pili are retracted, but the holdfast remains on the old pole from which the stalk emerges. In *Caulobacter crescentus*, the flagellum and pili are important for initial surface attachment while the holdfast is required for permanent attachment to a range solid surfaces including glass, mica, plastics, and decaying biotic material (23, 26). In fact, robust surface attachment via the holdfast adhesin is the characteristic that initially led to the isolation of *Caulobacter* species (27, 28). The holdfast also mediates polar cell-cell attachments resulting in the generation of multicellular structures, often called rosettes.

While the chemical composition of the holdfast material is not well understood, genetic and biochemical analyses indicate it is a polysaccharide (29, 30) that contains four major sugars (31, 32). There is also evidence that protein and DNA are important components of this adhesin (33). The role of the *C. crescentus* holdfast and other surface structures, including the flagellum and type IV pili, in colonization of the air-liquid interface has not been investigated.

In this study, I describe the process by which *C. crescentus* colonizes the air-liquid interface under static growth conditions, and define molecular determinants of this colonization process. Initially, cells accumulate as individual cells evenly dispersed in a monolayer at the air-liquid interface. At sufficiently high density, the monolayer transitions to a dense multilayered pellicle structure composed primarily of large connected rosette aggregates. Polar cell surface appendages including the flagellum, type IV pili and the holdfast all contribute to the development of this *C. crescentus* pellicle. As in biofilm formation on solid substrates, the flagellum and pili are important for efficient pellicle biofilm development, though neither is strictly required. Holdfast biosynthesis, on the other hand, is absolutely required for *C. crescentus* cells to accumulate at the air-liquid boundary and to form a pellicle. This work establishes a critical ecological role for the holdfast adhesin, namely in partitioning of cells to the air-liquid interface. Moreover, this work establishes the pellicle as a new system to study biofilm development in *C. crescentus* that is complementary to biofilm studies on solid surfaces.

## Results

### Caulobacter crescentus forms a pellicle under static growth conditions

To measure attachment to solid surfaces, bacteria are typically grown in polystyrene microtiter dishes or glass culture tubes, and surface attached bacteria are detected by staining with crystal violet. When grown in static culture (i.e. without shaking), *C. crescentus* cells accumulate in high numbers on glass or polystyrene near the air-liquid interface (see Figure 1b, bottom panel). This could reflect a bias in surface colonization at the boundary where the solid surface, culture medium, and air meet. Indeed, bacteria at this interface are reported to undergo rapid, irreversible attachment to solid surfaces at a level that is higher than cells in the bulk (10). However, it may also be the case that the enrichment of *C. crescentus* cells at the solid-liquid-air boundary simply reflects biased colonization of the entire air-liquid interface at the surface of the growth medium.

**Figure 1:**
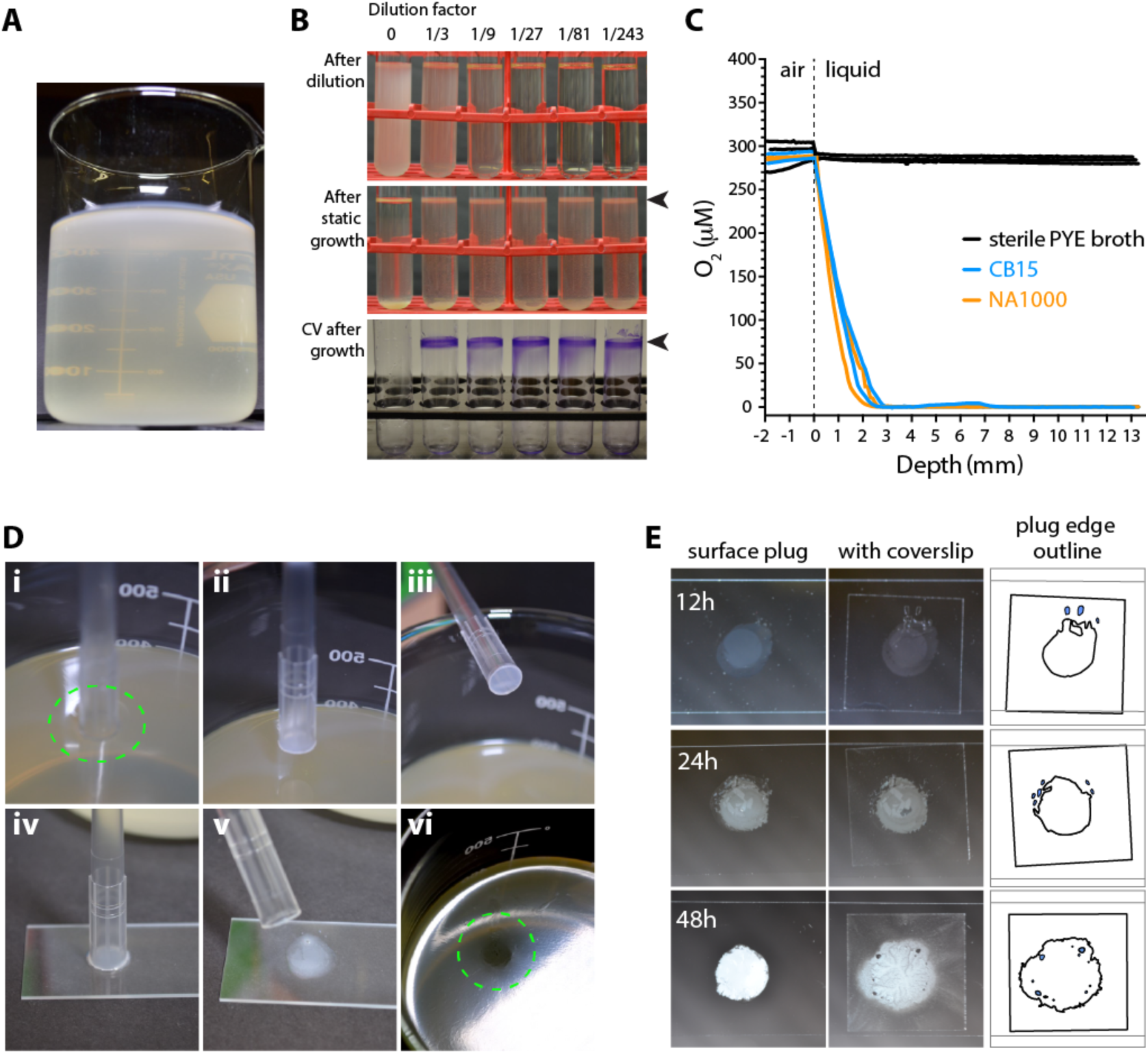
*Caulobacter crescentus* strain CB15 develops a pellicle at the air-liquid interface during static growth. **A.** Wild-type *C. crescentus* CB15 culture grown at room temperature without mixing (i.e. static growth) for three days. Note the accumulation of cells in a pellicle at the air-liquid interface at the top of the beaker. **B.** Pellicle development requires growth. *Top:* A culture was grown to stationary phase under aerated conditions, transferred to a fresh tube (far left) and serially diluted with fresh medium (towards the right; dilution fractions shown above each tube). *Middle:* The same tubes are shown after incubation on the benchtop for 4 days. Arrow highlights colonization of the air-liquid interface in diluted cultures, which grew post-dilution, but not in the undiluted culture. *Bottom:* Crystal violet (CV) stain of attached cells in tubes after cultures were washed away. The arrow highlights the position of the air-liquid interface. **C.** Oxygen gradient is steep at the surface of unmixed cultures. Oxygen concentration as a function of depth from the surface (0 mm) was measured in beakers in which PYE medium was left sterile (black traces), or inoculated with *C. crescentus* CB15 (blue traces) or NA1000 (orange traces) and incubated without mixing. CB15 cells accumulate at the air-liquid interface, while NA1000 cells are evenly dispersed throughout the culture. Both genotypes yield comparable oxygen gradients. Each trace represents an independent culture (n=2). Limit of detection is 0.3 μM. **D.** Method for sampling the pellicle. Large end of a sterile pipet tip is touched on the pellicle surface (i), lifted (ii, iii) and placed on a glass slide (iv, v). A pellicle scar (vi, green circle) can be seen after the plug removed from this 72-hour culture. **E.** Pellicle plugs were placed on glass slides (left) and then covered with a coverslip (middle). Outlines of the plugs under coverslips are on the right. Heavy line corresponds to the edges of the plugs. Stationary bubbles formed upon placement of the coverslip are filled in blue. The time since inoculation is indicated for each sample.

To visualize and monitor colonization of the air-liquid interface, I grew wild type *C. crescentus* strain CB15 statically in large volumes of a peptone-yeast extract (PYE) broth. A recent genome-scale analysis of *C. crescentus* indicates that complex media, such as PYE, are a more ecologically-relevant cultivation environment than a mineral defined medium such as M2 (34). In these conditions, as culture density increased, cells formed a surface film, or pellicle, that evenly covered the entire air-liquid interface (Figure 1a). Growth was required for pellicle formation: cultures grown to stationary phase in a roller or shaker did not form pellicles when transferred to static conditions unless diluted with fresh growth medium (Figure 1b). Static growth was accompanied by the establishment of a steep oxygen gradient in the culture flask. Dissolved oxygen levels were saturated in sterile growth medium across the measured depth of the culture flask. In contrast, oxygen was measurable in only the first 2-3 mm from the air-liquid interface in medium inoculated with cells. This was true for *C. crescentus* strains that develop pellicles (i.e. CB15) and strains that do not (i.e. NA1000) (Figure 1c).

Biofilm development on solid surfaces is a robust area of study in part due to the development of powerful methods to visualize live cells attached to glass slides in flow chambers (35) and to quantify cells attached to surfaces by crystal violet staining (36). Neither of these techniques is directly applicable to the study of biofilm pellicle development at the air-liquid interface. As such, I developed a method to image *C. crescentus* cells from the pellicle. An intact plug of the pellicle could be captured by using the large end of a 1 ml pipet tip (Figure 1d). This plug could be transferred to a glass slide and *a)* covered with a coverslip for visualization by light microscopy (Figure 1e), or *b)* allowed to adhere to the glass slide and stained with crystal violet (Figure 2). I used these techniques to monitor pellicle development in static cultures starting at low-density (OD_660_ ≈ 0.005).

**Figure 2:**
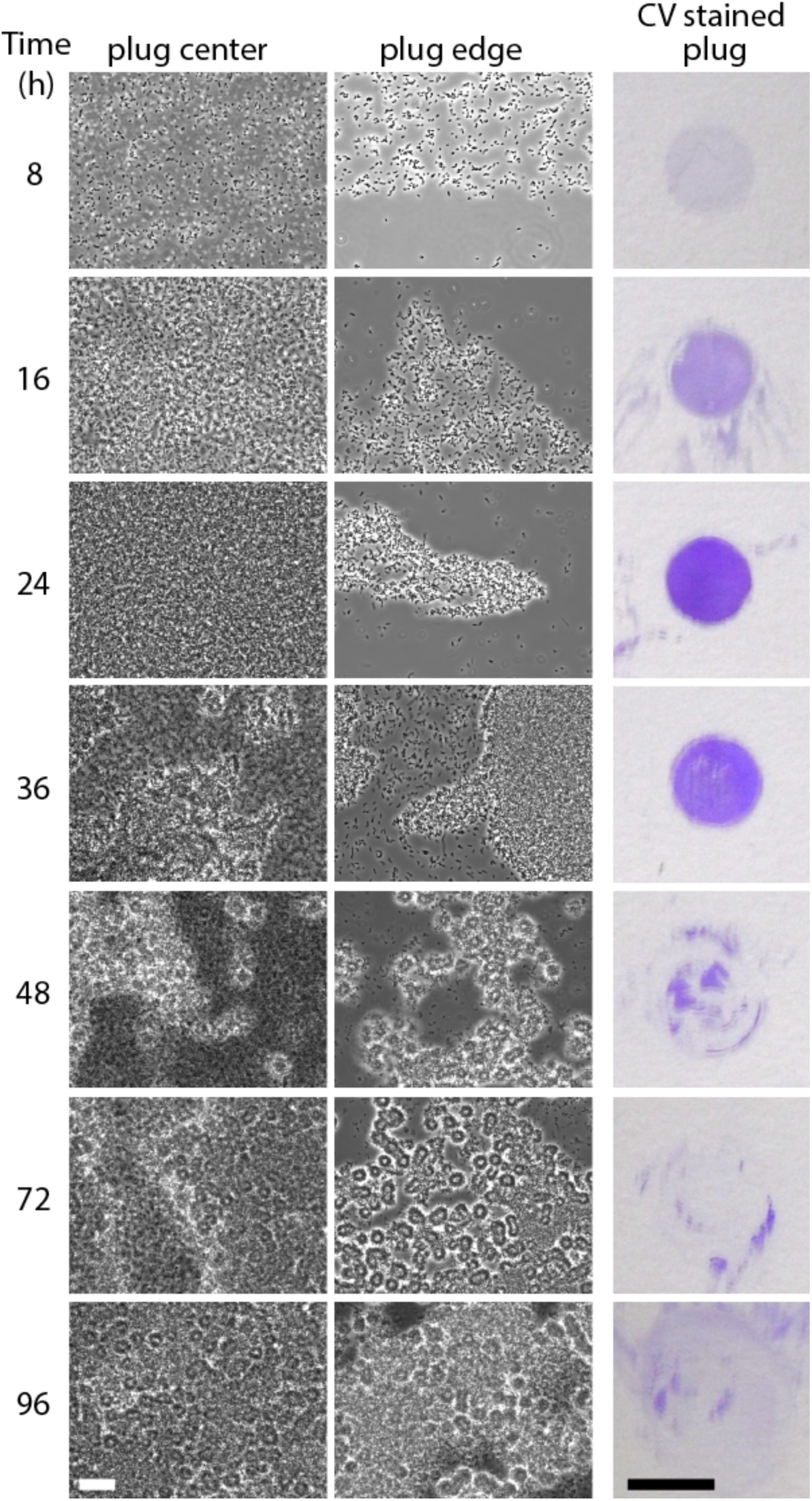
The pellicle develops from a homogeneous monolayer into a multilayered structure of dense rosettes. Surface plugs from a wild-type culture sampled periodically throughout static growth (time in hours after inoculation on the left) evaluated by phase contrast microscopy (left) and crystal violet (CV) staining (right). Two microscopy images are presented for each time point to capture the structure of cells in the center of the plug (left column of images) and also at the edges of the plug (right column; 8-36 hour samples) or cells disrupted from the multilayered plug structure (right column; 48-96 hour samples). Plug edges are outlined in Figure 1e. White scale bar is 20 μm. Black scale bar is 1 cm. This time course was repeated at least three times. Representative images from one experiment are presented.

Phase contrast imaging of plugs from the air-liquid interface revealed a rapid accumulation of cells at this boundary (Figure 2). Within hours, cells formed an evenly dispersed monolayer at the liquid surface. Through time, monolayer density increased and eventually formed a cohesive network of cells. Cell density in the surface layer was distinctly greater than in the sub-surface (bulk) medium (Supplemental Figure 1), which suggested that cells partition to the air-liquid interface. By 24 hours post inoculation, the surface monolayer had few, if any, gaps between cells. At this point, cells accumulated to a density at the surface of the liquid that was high enough to be visible as a film to the naked eye. In the monolayer stage, *C. crescentus* cells in plugs readily adsorbed to a glass surface and could be stained by crystal violet (Figure 2). These plugs maintained well-formed edges (Figure 1e and 2) and increased crystal violet staining of plugs was coincident with increased density of the monolayer. The void left by removing a plug from the surface film was rapidly filled by the surrounding film at this stage, suggesting that an early stage pellicle has fluid-like properties.

Between 24 and 48 hours, a transition occurred from a monolayer to a multilayered structure that contained dense rosettes (Figure 2). Simultaneously, the plug appeared less fluid and more cohesive, i.e. removal of a plug from the pellicle at this stage left a visible scar that was not filled by surrounding cells (Figure 1d). Upon this transition to a multilayered rosetted structure, pellicle plugs no longer adhered to a glass slide. Instead, the plugs crumbled and washed away during staining. These thick multilayered pellicle structures were challenging to image by light microscopy. When flattened by a glass coverslip, the structures were compressed and/or dispersed; regions of the plug that were less flattened by the coverslip appeared glassy when visualized by phase contrast. In either case, it is clear that the mature pellicle consists of a dense network of connected rosettes. These connections were often strong enough to maintain rosette interactions under the fluid flow that was induced by placing a coverslip on the pellicle plug (Supplemental Movie 1). Between 48 and 96 hours, the pellicle became even thicker and more visible macroscopically and microscopically. At some point after 96 hours, pellicles typically crashed, sinking under their own weight and settled in fragments at the bottom of the culture container.

### Holdfast are prominent in the pellicle

Many *Alphaproteobacteria*, including *C. crescentus*, form multicellular rosettes by adhering to each other through the polar polysaccharide, or holdfast. Given the notable presence of rosettes in the pellicle, I sought to directly visualize the holdfast in the pellicle using fluorescent wheat germ agglutinin (fWGA), a lectin that binds to N-acetylglucosamine moieties in the holdfast polysaccharide. Typical holdfast staining protocols using fWGA involve pelleting and washing the cells. To minimize disruptions to the pellicle structure during staining, I supplemented the medium with fWGA at the time of inoculation rather than staining after the pellicle was formed. I grew static cultures in the presence of 10, 1 or 0.2 ug/ml fWGA. The highest concentration delayed pellicle development (data not shown). Similarly, high concentrations (50 ug/ml) of WGA reduce holdfast adhesiveness to glass (37). In cultures with 1 or 0.2 ug/ml fWGA, pellicles developed similar to paired cultures without fWGA. I used 1 ug/ml of fWGA for these experiments, as signal was more intense than with 0.2 ug/ml.

In the early monolayer stages, nearly every cell was decorated with a holdfast at one cell pole (Figure 3). Fluorescent puncta corresponding to holdfast merged as the monolayer increased in density. As a multilayered structure emerged (32 hrs) distinct patterns of holdfast staining were evident in the different layers. The top layer (i.e. closest to air) contained a dense array of holdfast puncta similar to that observed in the monolayer at 24 hours. The lower layers of the plug contained a network of apparently inter-connected rosettes whose cores stained prominently with fWGA (Figure 3 (32 hr) and Figure 4). Bright fWGA puncta from rosette cores were observed in linear chains both in the center of intact pellicle plugs and in disrupted pellicle fragments (Figure 4 and Supplemental Figure 2). As the pellicle matured, the lower layers became packed with rosettes. The cores of adjacent rosettes were connected in three dimensions in a manner that likely confers strength to the pellicle biofilm.

**Figure 3:**
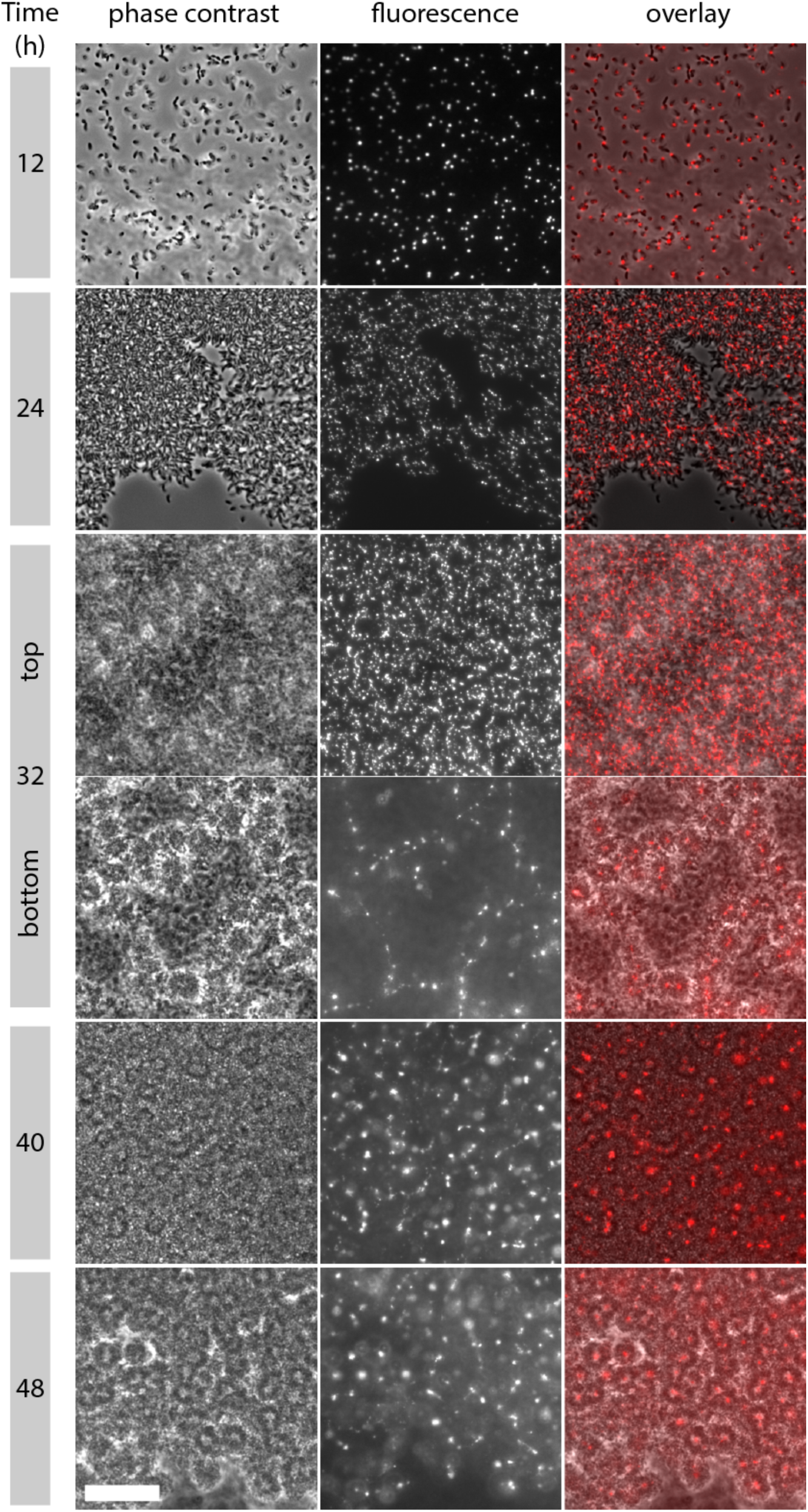
*In situ* fluorescent wheat germ agglutinin (fWGA) stained pellicle samples. Phase contrast and fluorescence images of cells grown in the presence of 1 ug/ml fWGA sampled at time intervals after inoculation. During the transition from a monolayer to a multilayer structure, at 32-hours, two focal planes of the same position in the pellicle plug are presented. These images correspond to the uppermost plane where fWGA bound to individual cells is in focus, and the bottom plane just below the monolayer where the centers of rosettes are in focus. At 40 and 48 hours, focal planes from the middle of the film are shown. Scale bar is 20 μm. Representative images from one of several timecourses are presented.

**Figure 4:**
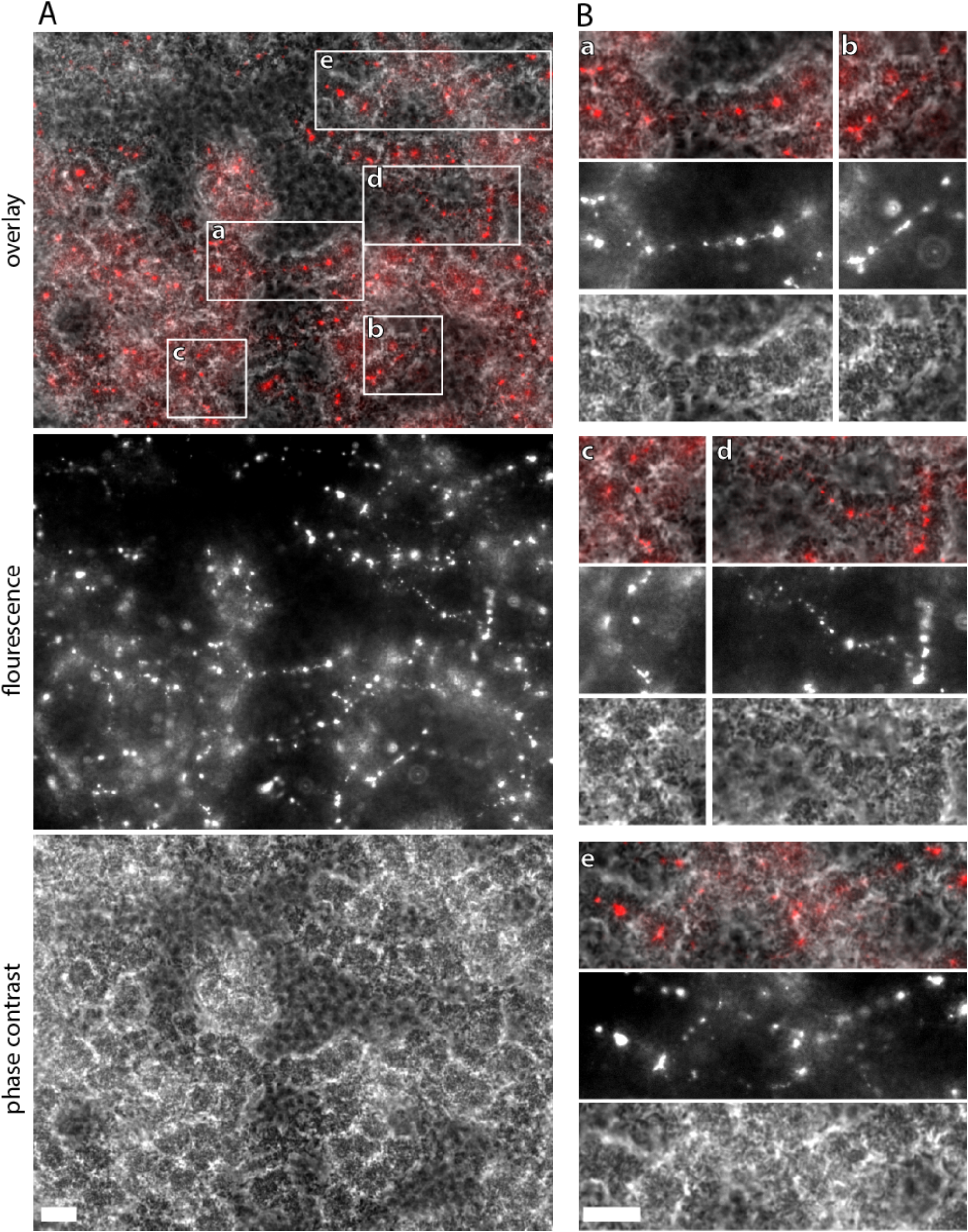
Linear arrays of rosettes in the center of an excised pellicle plug. **A.** In situ fluorescent wheat germ agglutinin (fWGA)-stained surface film harvested 40 hours after inoculation. The focal plane is just below a monolayer. Overlay (top), fWGA stained holdfast (middle) and phase contrast (bottom) images from one field of view (137 μm X 104 μm). Note, some of the rosette chains extend beyond the focal plane. **B.** Crops corresponding to the regions boxed in A. Scale bars are 10 μm.

In fragments of dispersed pellicle film, the spatial relationship between stained holdfast and the connected rosettes was more easily visualized (Supplemental Figure 2). Several types of structures are apparent. The tight focus of fWGA seen in radially-symmetric rosettes is consistent with holdfast adhered to each other at a single point. The cores of oblong rosettes are filled with many bright fWGA puncta and also a more diffuse fluorescent signal. This pattern suggests the rosette center is filled with holdfast material. The cores of each holdfast in these rosettes do not bind a singular central focus, but rather adhere in a mesh-like array.

### The long axes of cells are oriented perpendicular to air-liquid boundary, with the holdfast at the interface

Imaging cells in surface layer plugs provides evidence that the holdfast directly positions *C. crescentus* at the air-liquid interface. Small, static bubbles occasionally form in the process of mounting a pellicle plug on a glass slide (see Figure 1e). These bubbles present the opportunity to observe cells at high magnification at an air-liquid interface. The long axis of cells at this interface was perpendicular to the boundary between the air bubble and liquid, with the holdfast positioned directly at the interface (Figure 5). Moreover, when liquid flowed along stationary bubble edges, rafts of cells could be observed sliding along the boundary in the direction of flow (Supplemental Movies 2 and 3). Cells in these rafts were oriented perpendicular to the interface. The behavior of the cells attached to and moving along the bubble boundary was distinct from cells tumbling in the flow adjacent to the boundary.

**Figure 5:**
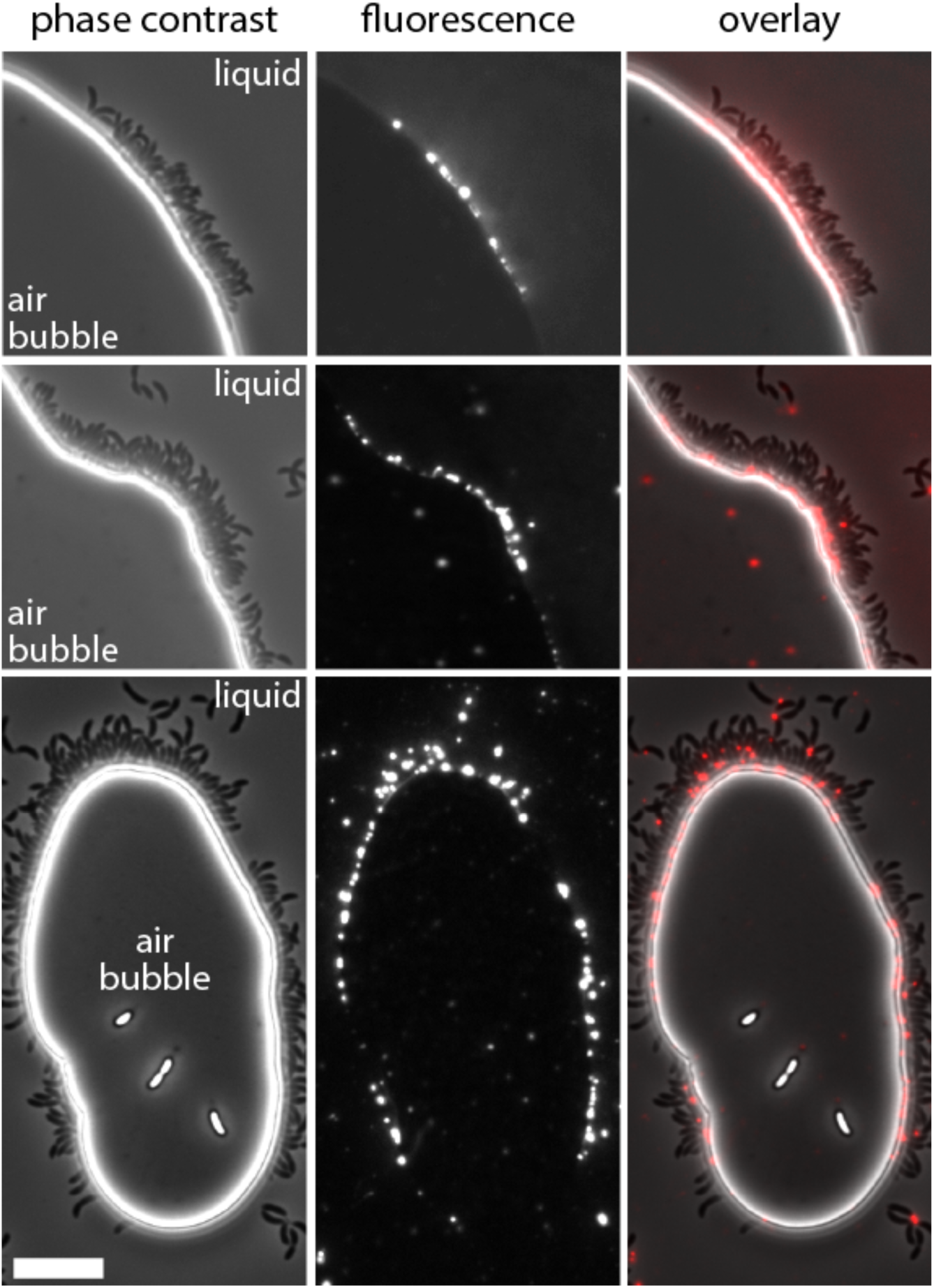
*C. crescentus* cells localized to the boundaries of air bubbles. Phase contrast (left), fluorescent wheat germ agglutinin (fWGA)-stained holdfast (middle) and overlay (right) micrographs from static cultures grown with fWGA, 8 to 12 hours after inoculation. The interface between the air bubbles and the liquid medium is bright in phase contrast. The air and liquid sides of the boundary are indicated. Scale bar is 10 μm.

### Holdfast biosynthesis is required for pellicle formation

Based on my observation of *1)* individual cells with holdfasts that occupy the air-liquid boundary and *2)* the presence of networks of rosettes in the pellicle, I tested if the holdfast is necessary for pellicle formation. Strains lacking *hfsJ*, a gene required for holdfast synthesis (38), do not form macroscopically visible pellicles (Figure 6). Not surprisingly, cells captured from the surface of Δ*hfsJ* cultures do not attach to glass slides as evidenced by the lack of crystal violet staining (Figure 7). At a microscopic scale, Δ*hfsJ* cells reach the surface microlayer as motile swarmer cells, but stalked and pre-divisional cells do not accumulate at the air-liquid interface (Figure 8). I obtained similar results for strains lacking a functional *hfsA* holdfast synthesis gene.

**Figure 6:**
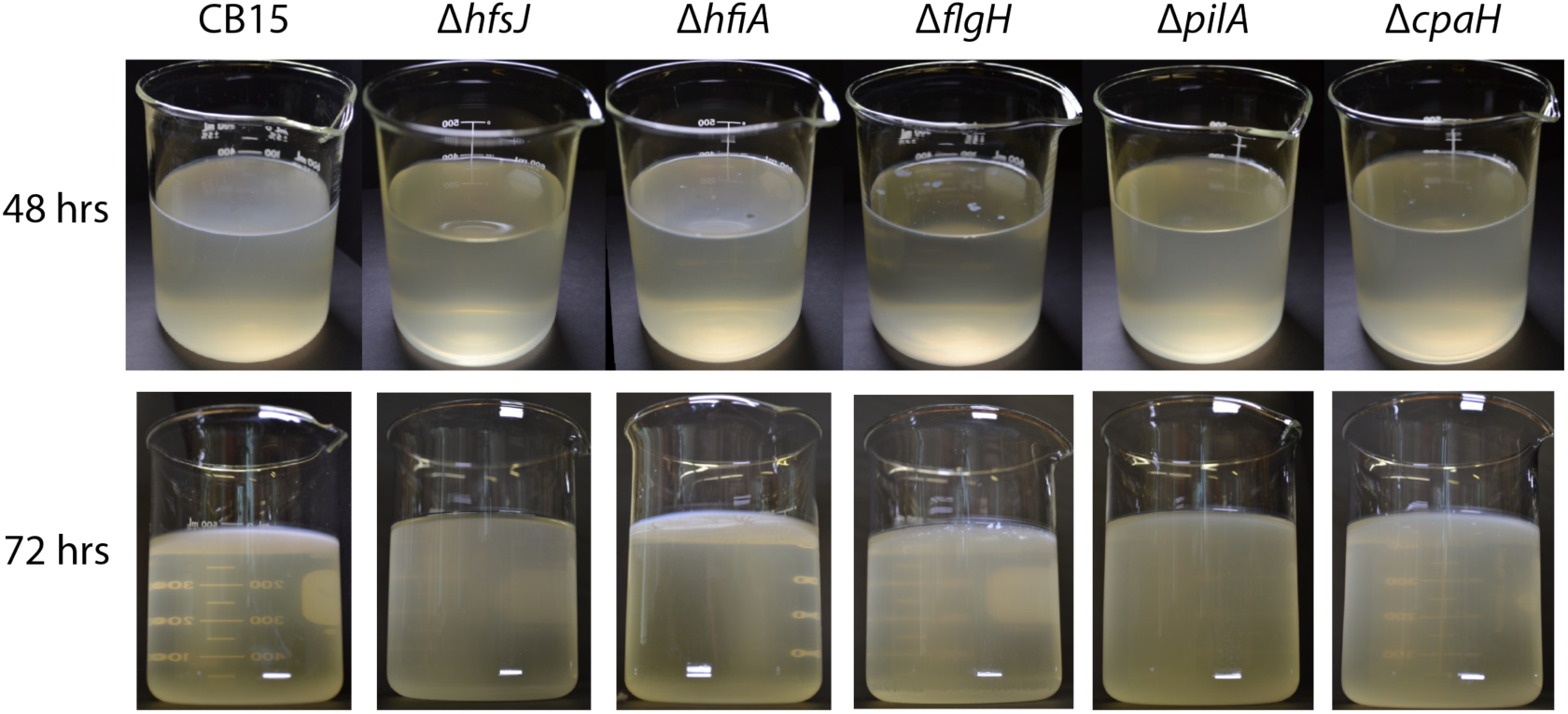
Macroscopic pellicles of polar appendage mutants. Static cultures of wild-type (CB15) and mutant (Δ) strains 48 and 72 hours after inoculation imaged from above or below respectively. See text for details on mutants. This experiment was repeated multiple times. One representative experiment is shown.

**Figure 7:**
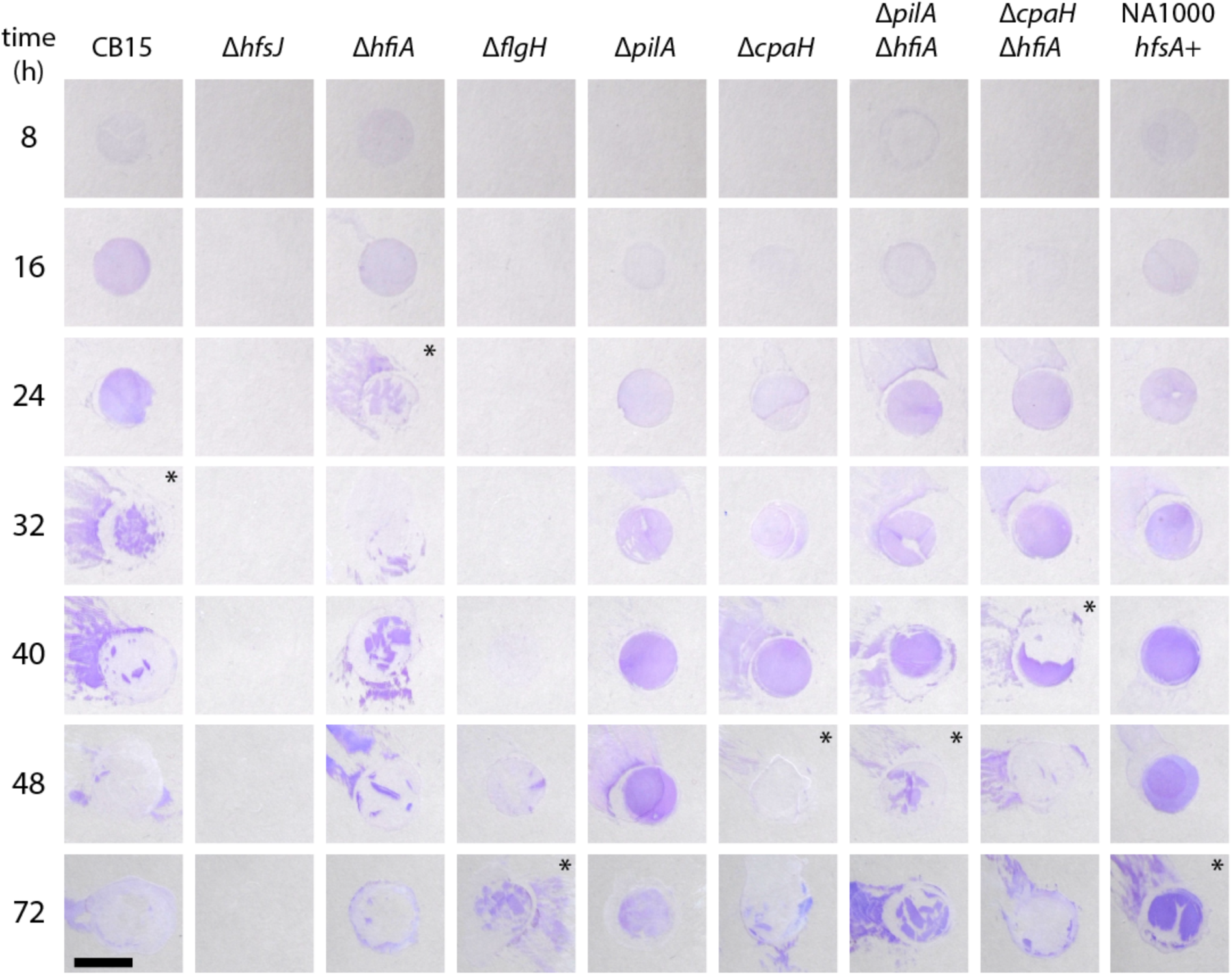
Crystal violet staining of pellicle plug samples. Pellicles of wild-type (CB15) and mutant (Δ) strains sampled throughout development and evaluated by crystal violet staining. Note three stages of pellicle development (CB15 times indicated): adhesive monolayer (up to 24 hours), crumbly transition phase (32-40 hours), and non-adhesive film (48+ hours). For each genotype, the beginning of the crumbly phase is marked with an asterisk. Pellicles sampled are from the same experiment presented in Figures 8. This experiment was repeated two additional times. Scale bar is 1 cm.

**Figure 8:**
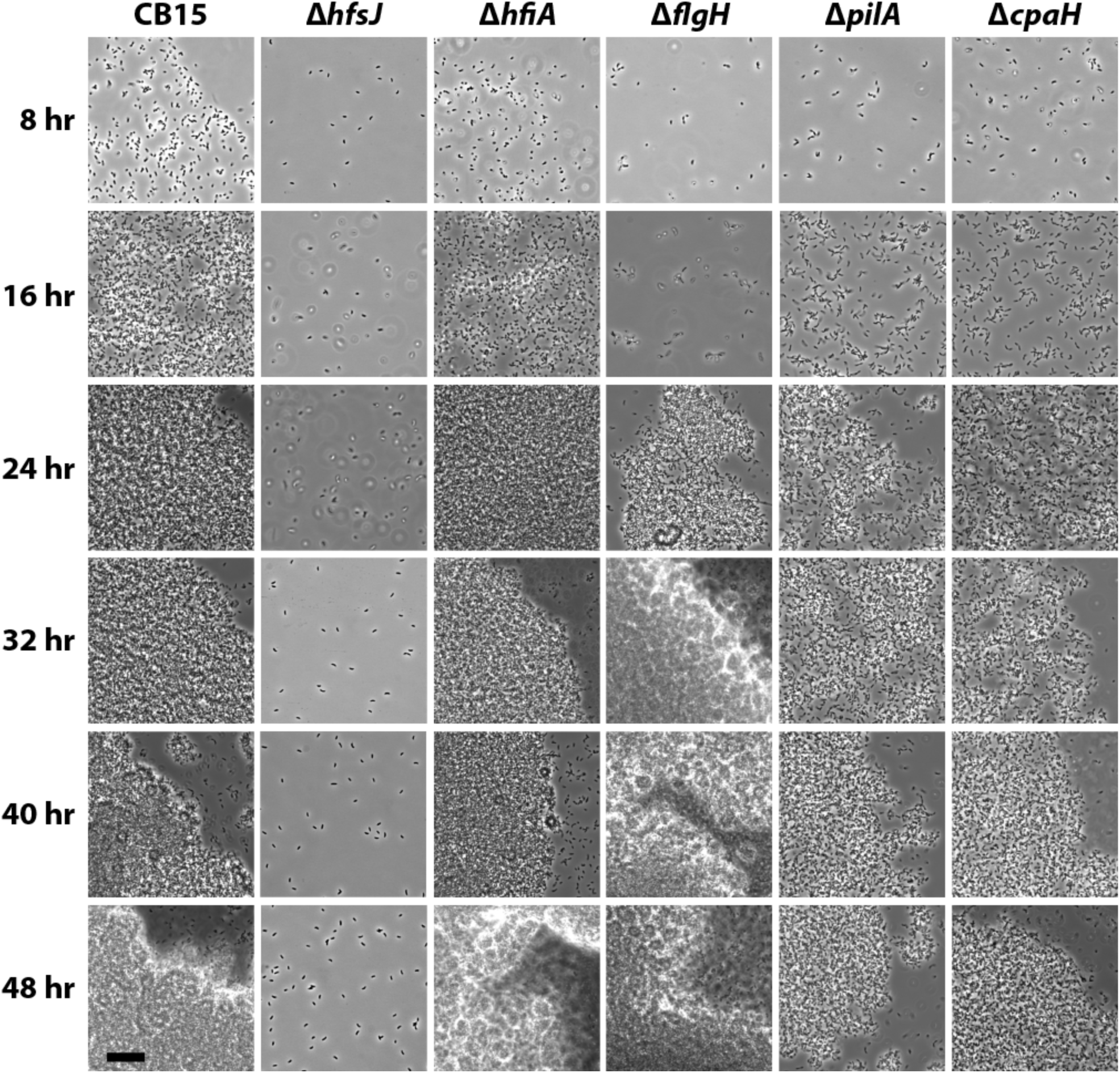
*C. crescentus* mutants lacking polar appendages exhibit defects in pellicle development. Phase contrast micrographs of pellicle samples from wild-type (CB15) and mutant strains as in Figure 2 sampled at 8-hour intervals. Scale bar is 20 μm. Representative images from one of three independent experiments are shown.

Holdfast biosynthesis is elevated in cells lacking *hfiA*, a negative regulator of holdfast biosynthesis (38). Pellicle development is accelerated in a Δ*hfiA* strain; these pellicles appear macroscopically thicker and leave plug scars at an earlier stage than wild type (Figure 6). Microscopically, the monolayer stage is similar to wild type (Figure 8), but the transition to a multilayered rosetted structure is more rapid, and the plugs lose adherence to glass sooner (Figure 7). Together these results indicate that holdfast is essential for cells to accumulate at the air-liquid interface and for the development of the pellicle structure. I conclude that enhancement of holdfast synthesis by deletion of *hfiA* promotes pellicle development.

### Flagella and pili determine efficient pellicle development

Flagella and pili are important factors for colonization of solid surfaces in *C. crescentus* (23, 26) and other species (39-41). Recently published data provide support for complex interplay between the flagellum, type IV pili and control of holdfast development in *C. crescentus* (25, 31, 42-45). Given the clear role of the pilus and flagellum in attachment to solid surfaces, and the regulatory connection between these structures and holdfast development, I tested the contribution of these appendages to *C. crescentus* pellicle development at the air-liquid interface. Specifically, I characterized pellicle development in a non-motile strain lacking *flgH*, which encodes the outer membrane ring component of the flagellum. In addition, I assessed the role of the type IV pilus in pellicle development using a mutant lacking *pilA*, which encodes the pilus filament protein, and a mutant lacking *cpaH*, which encodes a transmembrane component required for type IV pilus assembly.

Non-motile Δ*flgH* cells had dramatically delayed pellicle development. The pellicle that eventually emerged from this strain did not homogeneously cover the air-liquid interface, but rather contained microcolony-like aggregates that eventually became visible by eye at the surface of the culture (Figure 6). Δ*flgH* cells sampled at the air-liquid interface were primarily stalked or pre-divisional. At early time points, patches of cells attached to the coverslip, and small rosettes of 3-10 of cells were abundant. Small rosettes were rarely observed in the surface samples from other strains. Larger rosettes and aggregates were also evident in Δ*flgH* pellicles (Figure 8). With time, microcolonies consisting of dense mats of large rosettes became visible by eye (Figures 6), and when these large surface colonies were placed between a slide and a coverslip, clusters of rosettes became detached from the sample (Figure 8, see 40 hr sample). Eventually, the surface of the culture medium became covered with a film that did not adhere efficiently to glass and fragmented into large pieces (Figure 7, 72 hours). Though the non-motile Δ*flgH* strain was unable to actively move to the air-liquid interface, the hyper-holdfast phenotype of this strain (31, 44) seemed to enable capture of cells that arrived at the surface by chance. This resulted in cell accumulation and formation of the observed microcolonies at this boundary. I postulate that the inability of Δ*flgH* daughter cells to disperse, combined with premature holdfast development in this strain (44) promotes microcolony formation rather than a uniform distribution of cells at the air-liquid interface. These data support a model in which flagellar motility enables cells to efficiently reach the air-liquid interface, but that motility *per se* is not required for cells to colonize this microenvironment.

Both Δ*pilA* and Δ*cpaH* strains were defective in pellicle development. These pilus mutants are motile and capable of synthesizing holdfast. Both mutants accumulated at the air-liquid interface as monolayers similar to wild type (Figures 8). However, the density of these monolayers increased more slowly than wild type. In addition, surface plugs from these mutant films retained the capacity to adhere to glass for a longer period (Figure 7), and resisted scaring upon plug removal for an extended period of sampling. These observations are consistent with an extended monolayer phase. Even when dense monolayers formed, both mutants were defective in transitioning to a multilayered structure as evidenced by microscopic images and crystal violet stains of surface plugs (Figures 7 & 8).

It is notable that in a selection for mutants with surface attachment defects, these two mutants displayed distinct phenotypes; Δ*pilA* mutants had reduced surface attachment, while Δ*cpaH* mutants displayed enhanced surface attachment owing to increased holdfast synthesis (31). Thus in the context of attachment to solid surfaces, increased holdfast synthesis can outweigh defects from the loss of pili. In pellicle development on the other hand, the defects in these two classes of pilus mutants were nearly the same. The primary difference was that the Δ*cpaH* mutant transitioned to a non-adherent, crumbly film sooner than the *ΔpilA* mutant, as might be expected for a strain with elevated holdfast synthesis (Figure 7). Even though *ΔcpaH* transitioned to a crumbly structure sooner than *ΔpilA*, this mutant was still significantly delayed compared to wild type. In addition, micro-colonies were often observed in *ΔcpaH* surface films, but were smaller and less pronounced than in the Δ*flgH* surface films (Figure 6).

Finally, I examined pellicle development in Δ*pilA* and Δ*cpaH* mutants that also carried an in-frame deletion of *hfiA* in order to test whether elevated holdfast production could overcome the defects associated with the loss of pili. The Δ*pilA*Δ*hfiA* and Δ*cpaH*Δ*hfiA* double mutant strains did transition to a crumbly, non-adherent film sooner than their Δ*pilA* and Δ*cpaH* counterparts. However, both double mutant strains were still delayed compared to the Δ*hfiA* single mutant and were not restored to wild-type pellicle development (Figure 7). Together, these data indicate that pili are not required for *C. crescentus* to colonize the air liquid interface, but these appendages do contribute to formation of a dense, robust pellicle. Moreover, these data indicate that elevated holdfast production promotes pellicle development, but is not sufficient to fully compensate for the loss of pili.

### Pellicle architecture is influenced by a mobile genetic element

NA1000 is a standard laboratory strain that is almost completely isogenic with CB15 (46), and that is used to produce synchronized populations of *C. crescentus* for cell cycle studies (47). The synthesis of an extracellular polysaccharide (EPS) on the surface of stalked cells, but not newborn swarmer cells, enables isolation of NA1000 swarmer cells by centrifugation in percoll (48). Genes required for the synthesis of this cell-cycle regulated EPS are encoded by a mobile genetic element (MGE) that is present in the NA1000 genome, but missing from CB15 (46). In addition, NA1000 is defective in holdfast formation owing to a frame-shift mutation in *hfsA* (46).

NA1000 did not develop pellicles under static growth conditions. Restoration of *hfsA* to a functional (non-frameshifted) allele was sufficient to enable pellicle formation in this background (Figure 9a). However, NA1000 *hfsA*^+^ pellicles were qualitatively different from CB15 in many respects. NA1000 *hfsA*^+^ pellicles were more fluid, i.e. voids from pellicle plugs quickly filled in rather than leaving scars. In addition, plugs from mature NA1000 *hfsA*^+^ pellicles did not crumble like CB15 (Figure 7). At a microscopic level, I observed more space between cells in the center of the film and in dispersed rosettes. NA1000 *hfsA*^+^ rosettes were less tightly packed and more interwoven (Figure 9b). In short, even though restoration of the *hfsA* frameshift in NA1000 restores holdfast development and pellicle formation, there are significant phenotypic differences in cell packing and pellicle architecture between *C. crescentus* strains NA1000 and CB15.

**Figure 9:**
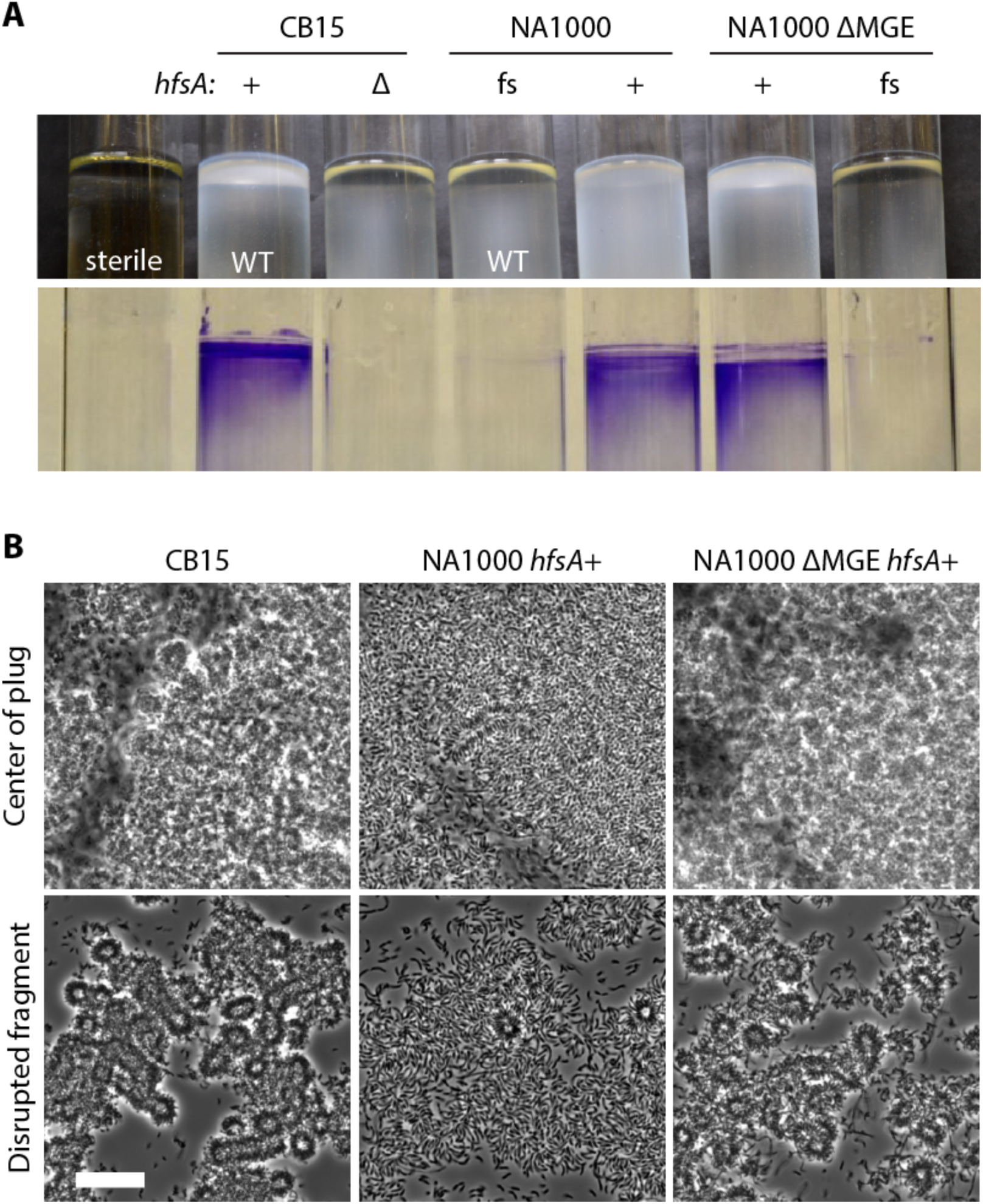
Pellicle structures of NA1000 strains differ from CB15. **A.** Pellicles of cultures grown statically for 3 days are pictured (top). After growth, the tubes were stained with crystal violet to highlight cells adhered to the glass at the surface of the cultures (bottom). Genotypes of strains are indicated above the tubes. Strains that differ only at the *hfsA* locus are paired and the *hfsA* allele is indicated: functional CB15 allele (+), null (Δ), or frame-shifted NA1000 allele (fs). See text for details about the mobile genetic element (MGE). A tube with sterile medium (left) demonstrates the characteristics of an uncolonized meniscus, similar to what is seen at the surface of medium colonized with strains lacking a functional *hfsA* allele. The whole surface of the culture is opaque when a pellicle film is present. **B.** Phase contrast micrographs of pellicle samples from CB15, NA1000 *hfsA*+ and NA1000 ΔMGE *hfsA*+ pellicles collected 48 hours after inoculation. The center of the plug (top) and rosettes disrupted from the film (bottom) were imaged. Scale bar is 20 μm.

Based on these observations, I reasoned that the NA1000-specific EPS (synthesized by gene products of the MGE) may be responsible for pellicle differences between CB15 and NA1000. To test this hypothesis, I grew static cultures of an NA1000 isolate from which the MGE had spontaneously excised (NA1000 ΔMGE) (46) and an isogenic strain in which I restored the null frameshift mutation in *hfsA* (NA1000 ΔMGE *hfsA*^*+*^). As expected, the NA1000 ΔMGE strain with the frameshifted *hfsA* allele did not accumulate at the air-liquid interface, and restoration of *hfsA* enabled pellicle development (Figure 9a). Loss of the MGE, and thus cell-cycle regulated EPS, resulted in a pellicle architecture that more closely resembled CB15 with closer cell packing and more compact rosettes (Figure 9b). In addition, the enhanced buoyancy conferred by the MGE was apparent when observing culture medium below the surface. CB15 and NA1000 ΔMGE *hfsA*^+^ cells not trapped at the air interface tend to settle to the bottom. NA1000 *hfsA*^+^ cells were more evenly distributed throughout the depth of the culture (Figure 9a), presumably owing to the cell-cycle regulated EPS present on the non-motile stalked cells (46, 48).

It is worth noting that several phenotypic differences between CB15 and NA1000 were not determined by the presence/absence of the MGE. Specifically, NA1000 derived cells were notably larger than CB15 and more prone to filamentation in the pellicle context, regardless of the MGE. The genetic polymorphisms responsible for these phenotypic differences have not been determined.

The MGE present in NA1000 strains accounts for the major differences in cell packing and pellicle architecture between CB15 and NA1000. This observation supports a model in which modulation of secreted polysaccharides have profound effects on cell-to-cell interactions, and possibly on cell-to-interface interactions at boundaries. These strain differences should be considered as investigators look toward future studies of *C. crescentus* attachment behavior and biofilm development.

### *Caulobacter* accumulates at the air-liquid interface in dilute media

The thick pellicle films observed in static culture in PYE medium involve high densities of cells; PYE can support growth to 10^8^-10^9^ CFU/ml. I wondered if *C. crescentus* could form a pellicle film in a more nutrient limited environment, i.e. in a medium that did not support such high cell density. To address this question, I used a series of increasingly dilute complex media rather than a mineral defined M2-based medium for two main reasons: 1) standard M2-based media support dense cell growth (i.e. are not growth limiting), 2) there is evidence that complex media better reflect the complex suite of nutrients encountered by *Caulobacter* in natural ecosystems (34). While progressive dilution of complex medium certainly reduced culture density (and cell size), a thick pellicle was observed in 0.1X PYE, large rosettes accumulate at the surface of 0.03X PYE, and small clusters of 2-4 cell rosettes form on the surface of cultures grown in 0.01X PYE (Figure 10). The density of cells in the pellicle was proportional to the carrying capacity of the growth medium. Importantly, in all these conditions, the distribution of cells is strongly biased toward accumulation at the air-interface (Figure 10). Thus even highly diluted medium, *C. crescentus* partitions to the air-liquid interface.

**Figure 10:**
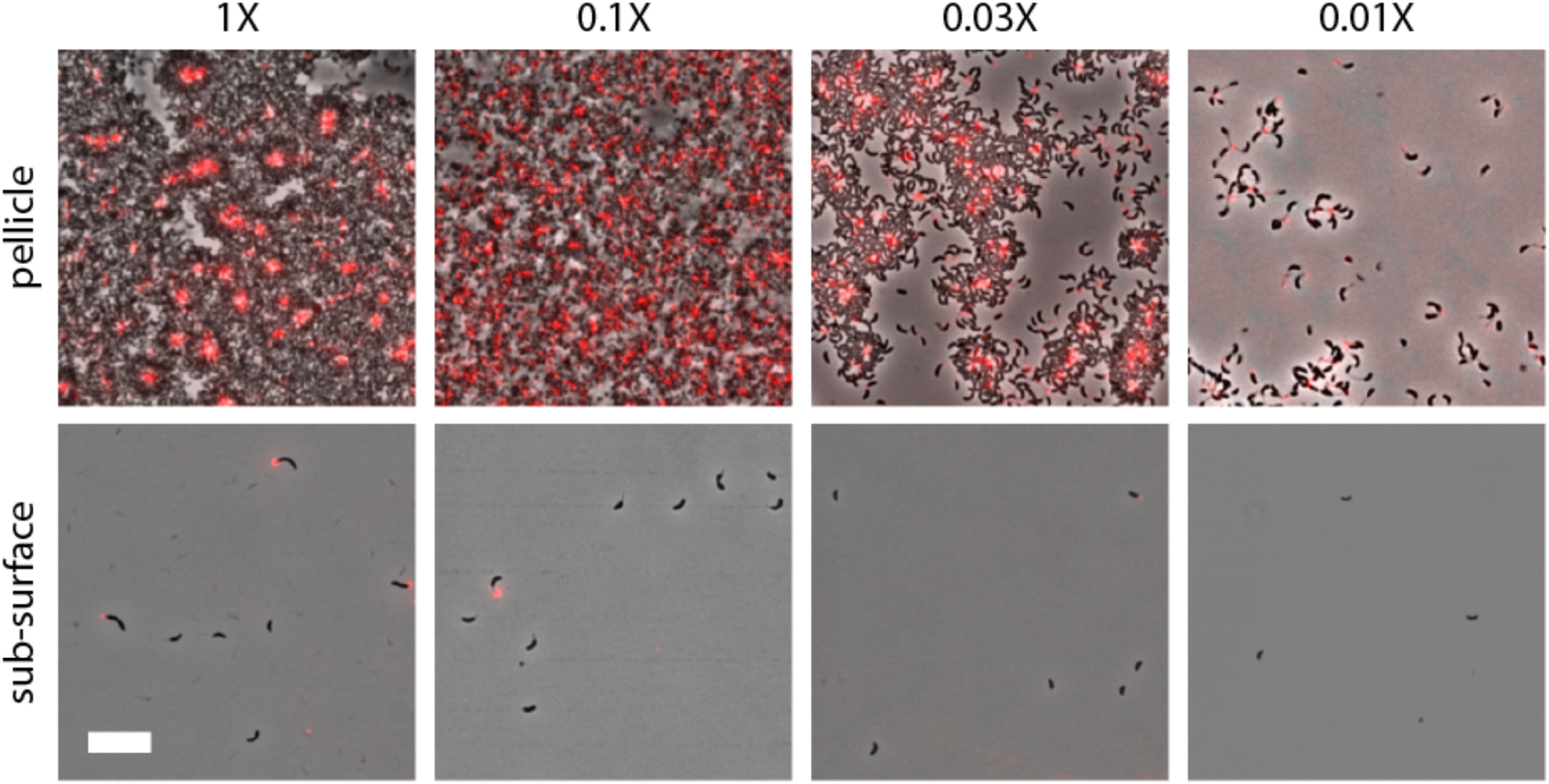
Cells partition to the air-liquid boundary in dilute complex medium. Surface and sub-surface samples of *C. crescentus* CB15 cultured in increasingly dilute PYE medium. Dilution factors from the standard lab recipe (see Materials and Methods) are indicated at the top. Cultures were grown statically with fWGA to stain holdfast for 2.5 days. Samples were imaged using phase contrast and fluorescence, and an overlay is presented for each condition. Scale bar is 10 μm.

## Discussion

### An alphaproteobacterial model for biofilm development at the air-liquid boundary

Molecular factors that contribute to colonization of solid surfaces in both environmental and host-microbe contexts are well understood for many bacterial species. While biofilms at the air-liquid boundary have been studied, they have received less attention and our understanding of the molecular determinants of biofilm development at such interfaces is less well developed. Data presented in this study define distinct stages of pellicle development in *C. crescentus*, a model *Alphaproteobacterium*. The *C. crescentus* pellicle does not initiate at the solid edges of the air-liquid interface, but rather develops uniformly across the entire liquid surface. Initially, individual cells accumulate at this boundary as an evenly dispersed monolayer of individual cells trapped at the interface. When the monolayer becomes sufficiently dense, rosettes accumulate beneath the monolayer and eventually form a multilayered pellicle structure comprised primarily of large dense rosettes (Figure 2). These stages are reminiscent of biofilm development on solid substrates, in which surfaces are often initially colonized with a monolayer of cells before more complex three-dimensional structures form. I propose that *C. crescentus* colonization and pellicle formation at the air-liquid boundary is an experimentally tractable model for the study of biofilm development in an *Alphaproteobacterium*.

While it is known that bacteria will form monolayers at air-liquid interfaces in natural settings, and I observed partitioning of *C. crescentus* to the air interface in 100-fold diluted PYE (Figure 10), the ecological relevance of the thick three-dimensional structures I observe in later stages of *C. crescentus* pellicle development is not clear. Poindexter describes individual prosthecate cells, but not rosettes in environmental samples; rosettes were only evident in her pure cultures (20). Similarly, in surface samples collected directly from a freshwater pond, Fuerst describes prosthecate cells, but does not note rosettes in this study (49). I am not aware of any descriptions of *Caulobacter*, or other rosette-forming *Alphaproteobacteria* producing rosettes outside of the laboratory. One expects that rosettes and three-dimensional *Caulobacter* pellicles/films should only occur in environments with sufficient nutrients to support high cell densities. That said, recent metagenomic analyses challenge the historical view that *Caulobacter* species are oligotrophs that exclusively inhabit dilute aquatic environments (18). *Caulobacter* species are broadly distributed in soil and aquatic environments and are most prevalent in systems with abundant decaying plant matter, which are not associated with nutrient limitation (18). Thus it may be the case that *Caulobacter* spp. do form thick pellicles in static aquatic systems rich in decaying plant material. Regardless, the holdfast clearly enables exploitation of the air-liquid interface, even at low cell density (Figure 10).

### Multiple polar appendages contribute to pellicle development

*C. crescentus* swarmer cells are born with a single flagellum and multiple pili that decorate the old cell pole, and are preloaded with the machinery to elaborate a holdfast at this same pole (20, 23, 24). Development of these surface appendages is intimately tied to the cell cycle and is central to the lifestyle and ecology of *Caulobacter* species. Specifically, the flagellum confers motility and enables swarmer cells to disperse, while the flagellum and the pili together contribute to reversible attachment during colonization of solid surfaces. When deployed, the holdfast confers irreversible attachment to solid surfaces (20, 23, 26, 37, 50). In colonization of the air-liquid interface, each of these appendages also plays important roles. Cells lacking a functional flagellum are unable to efficiently reach the interface and, instead, arrive there only by chance. Cells unable to synthesize holdfast reach the liquid surface as motile swarmers, but do not remain after differentiation into a non-motile stalked cell. Thus, holdfast mutants do not accumulate at the surface and cannot form a dense pellicle film. Finally, cells lacking pili efficiently reach the air-liquid interface and accumulate to high densities, but exhibit developmental delays. A synthesis and discussion of published data on the *C. crescentus* flagellum, pili, and holdfast in the context of my results follow.

#### FLAGELLUM

The requirement that *C. crescentus* be motile to efficiently reach the air-liquid interface (Figures 6 and 8) is not particularly surprising. Genes involved in aerotaxis and motility are known determinants of pellicle formation in other aerobes (eg. (16, 51-54)). *C. crescentus* is capable of aerotaxis (55), though the requirement for aerotaxis *per se* in *C. crescentus* pellicle formation remains undefined as the sensors are unknown. While static *C. crescentus* cultures have a steep oxygen gradient at the air interface (Figure 1c), non-motile or non-adhesive *C. crescentus* mutants can still grow to high density in static culture. This is consistent with a tolerance of this species for microoxic conditions (56).

While motility is required for cells to efficiently reach the surface, I have shown that it is not explicitly required for accumulation at the surface. *ΔflgH* mutants, which lack a flagellum, colonize the air-liquid interface, albeit inefficiently and in a less uniform manner than wild type (Figures 6 and 8). It is known that the loss of flagellar genes including *flgH* and *flgE* results in a hyper-holdfast phenotype (31, 44). In the context of pellicle development, the observed microcolonies of rosettes in the *ΔflgH* strain suggests that its hyper-holdfast phenotype can overcome the motility defect of this strain.

#### HOLDFAST

Data presented in this study provide evidence that the holdfast can function to trap cells at the air-liquid interface. Mutants defective in holdfast synthesis cannot partition to the surface layer (Figures 6 and 8). Inspection of bubble surfaces formed when mounting samples for microscopy reveals cells positioned perpendicular to the bubble boundary with the holdfast pole occupying the air-liquid interface (Figure 5). I infer that the holdfast allows polar attachment of replicative stalked cells to air-liquid interfaces, similar to solid surfaces. The observations reported here are reminiscent of an earlier report that the Alphaproteobacterium *Hyphomicrobium vulgaris* stands perpendicular to air-liquid, liquid-liquid and solid-liquid boundaries with the replicative pole at the interface (57).

How might the holdfast enable cells to remain at this interface, and what can be inferred about the nature of the holdfast material from these observations? The microlayer between the bulk liquid and the air represents a unique physiochemical environment. Hydrophobic and amphipathic molecules partition to this boundary (1, 2, 7, 10). Surface hydrophobicity is an important feature of bacteria that colonize the air-liquid interface (58). Though the exact chemical nature of the holdfast is not known, the fact that it apparently partitions to this zone implies that it has hydrophobic, or at least amphipathic properties. A similar conclusion was reached regarding the unipolar polysaccharides secreted by the *H. vulgaris* and the unrelated Sphingobacterium *Flexibacter aurantiacus* (57).

The air-liquid interface of complex aqueous broth is more viscous than the bulk solution owing to polymers adsorbed at this surface (7, 8, 15, 59). Increased surface viscosity is responsible for trapping motile swarmer cells at the air-liquid interface (59) and may also trap the holdfast, which itself is secreted as an amorphous viscous liquid (60). In sum, the holdfast can apparently function to partition non-motile replicative cells to the air-liquid interface. This function is likely important for an aerobe that is only motile (and aerotactic) in the non-replicative swarmer phase of its life cycle.

How then do rosettes, in which the holdfast is buried in the interior of a cluster of cells, partition to the air-liquid boundary? The answer to this question is not clear from the data presented in this manuscript. One possibility is that the holdfast polymer excludes water from the rosette core to an extent that it reduces the density of the collective aggregate. More extensive biophysical characterization of rosettes will lead to a better understanding of the role of these structures in partitioning to the air-liquid interface and in pellicle development.

#### PILUS

Type IV pili are not required for cells to reach or adsorb to the air-liquid interface (Figure 8). However, cells lacking pili inefficiently reach high densities at the interface and are extremely delayed in the transition to a multilayered pellicle structure, even when holdfast production is elevated. I envision two non-exclusive explanations for this result: *a)* pili are important factors mediating cell-cell interactions and facilitate the coalescence of cells during rosette formation; *b)* pili constitute a matrix component that confers strength and rigidity to the pellicle. Pili can extend up to 4 um in length (61) and physically retract (43). Pilus interactions between neighboring cells should increase load during pilus retraction, thereby stimulating holdfast production (43) while simultaneously bringing holdfast bearing cell poles in closer proximity. In this way pili may organize cells and promote rosette development. This model is similar to *Neisseria gonorrhoeae*, where pilus interactions and pilus motor activity promote dense packing of cells (62-64). Electron micrographs of rosettes of the closely related species, *Asticcacaulis biprosthecum*, reveal a network of pili surrounded by holdfast at the junction between poles (65). These snapshots lead one to speculate that pilus retraction brought these cell poles together. In addition, the *A. biprosthecum* micrographs, combined with the results described here, inform the hypothesis that pili confer structural support to reinforce holdfast-mediated interactions between cells.

Although difficult to capture by standard light microscopy, I observed that assemblies of cells were less organized at the air-liquid interface in both pilus null strains (Δ*pilA* and Δ*cpaH*). In these mutants, it was often difficult to assess whether cells were arranged in a rosette (i.e. attached at the distal end of the stalked poles) or simply in an unordered clump of non-specifically adherent cells. This qualitative conclusion held true in analyses of pellicle plugs where I blinded the strain genotype. My observations support a role for the pilus in organizing and promoting cell-cell interactions. In many species, type IV pili mediate motility, however in *Caulobacter* the primary role of these appendages seems to be attachment to surfaces (26, 43, 66, 67). As an extension, I propose that *C. crescentus* pili facilitate cell-cell attachments in the context of the pellicle. The role of type IV pili in cell-cell interactions and robust pellicle formation merits further study.

Finally, I note that the role of pili in mediating attachment is context dependent. In a pellicle, mutants lacking the pilus filament *(ΔpilA*) or a component of the pilus assembly machine (*ΔcpaH*) exhibit similar phenotypes. In the context of attachment to cheesecloth or polystyrene, Δ*pilA* has attenuated surface attachment while a *ΔcpaH* strain exhibits hyper-attachment (31). In shaken broth, deletion of *cpaH* increases the fraction of cells with a holdfast while deletion of *pilA* does not affect the probability of holdfast development (31). On an agarose pad, cells lacking *pilA* exhibit delayed holdfast development (44). Collectively, these results indicate that physical/environmental constraints likely influence the relative importance of the pilus function *per se* and pilus regulation of holdfast development on attachment.

### On the formation of cell chains and the putative threads that connect them

Fluorescence imaging of pellicles reveals rosette cores as well as looser assemblies of cells that appear to be connected in linear arrays (Figures 3, 4 and Supplemental Figure 2). Properties of the rosette exterior may facilitate connections between rosettes, but rosette surface connections alone should be non-directional and result in randomly organized aggregations of rosettes. The linear nature of the connections suggests the possibility of a thread-like structure that does not stain with fWGA, but upon which holdfast-bearing cells can attach (Figures 3, 4 and Supplemental Figure 2). What, then, might be this material to which holdfasts adhere that could mediate longer-range interactions in a pellicle? The length of the connections suggests a polymeric molecule (polysaccharide, DNA or a protein fiber). This putative material does not bind WGA, suggesting it is not holdfast polysaccharide, unless the cell produces a modified form lacking N-acetylglucosamine. In the pellicle context, *C. crescentus* may synthesize a previously uncharacterized extracellular polysaccharide. For example, *Agrobacterium* elaborates a polar adhesin and also synthesizes extracellular cellulose fibrils that aid in cell aggregations and attachment to plant cells (68, 69). It is also possible that these threads are DNA. This molecule is a well-established component of the biofilm matrix of other bacteria (17). DNA associates with the outer layers of the *C. crescentus* holdfast and similarly is observed adjacent to the holdfast polysaccharide in rosette cores (33). In other work, DNA released during cell death was demonstrated to bind to holdfast and inhibit attachment (70). This suggests a model in which DNA associates with the holdfast polysaccharide and at sufficiently high concentrations masks the adhesin, similar to high concentrations of WGA (37). Finally, polymers of proteins such as pili or flagella could conceivably facilitate long-range interactions. Pili are observed in rosettes of *Asticcacaulis* (65) and cells lacking PilA are defective in development of these multicellular structures in the pellicle (Figure 8). However, this filament is typically retracted into the cell and single pilus filaments are too short to facilitate interactions of the length scale I observe. Flagellar polymers on the other hand are shed into the medium (71), though they are occasionally observed still attached at the end of a stalk extension (20). It may be the case that overlapped mixtures of these filamentous materials produce these ‘threads’. Future work will be necessary to identify this putative component of the *Caulobacter* pellicle biofilm.

### The distribution and ecological importance of the holdfast in Alphaproteobacteria

Synthesis of a holdfast-like adhesin at one cell pole is a broadly conserved trait in *Alphaproteobacteria*. Examples of species that secrete polar adhesins or form polar rosette aggregates have been described in almost every *Alphaproteobacterial* order including *Rhizobiales* (72-81), *Caulobacterales* (20, 37, 65, 82-84), *Rhodobacterales* (85-90), and *Sphingomonadales* (91-93). Exceptions are *Rhodospirillales* and *Rickettsiales*, which are at the base of the *Alphaproteobacterial* tree (94). The ensemble of holdfast synthesis genes (29, 30, 72, 79, 82, 88, 95), and chemical composition of the holdfast polysaccharides (37) vary between species and families, which may reflect chemical differences in the niches particular species colonize.

For many *Alphaproteobacteria*, the advantage of a polar adhesin for attachment to surfaces is obvious: *Agrobacterium* and *Rhizobium* adhere to plant roots during symbiosis, *Roseobacter* interact with algae in a symbiosis, and to submerged abiotic surfaces that are coated by conditioning films. I propose that attachment/partitioning to air-liquid interfaces is a general function of holdfast-like polar polysaccharides in some species. For example, *Phaeobacter* strain 27-4 and other *Roseobacter* spp. form interlocking rosettes at the air liquid interface in static cultures (86, 87). In biofilm assays, *Caulobacter* and *Agrobacterium* attach most robustly at the air-solid-liquid interface and this attachment requires a polar adhesin (eg. (44, 95) and Figure 1b). For *Alphaproteobacteria* that are aerobic heterotrophs, the advantage of a cellular mechanism to exploit elevated nutrients and oxygen at the air-liquid interface is clear. The holdfast can provide this function.

## Materials and Methods

### Growth conditions

The *C. crescentus* strains used in this study are derived from the CB15 wild-type parent unless noted; see Table 1. All strains were cultured in peptone-yeast extract (PYE) broth containing 0.2% peptone, 0.1% yeast extract, 1 mM MgSO_4_, 0.5 mM CaCl_2_. PYE was solidified with 1.5% agar for propagation of strains. Strains detailed in Table 1 were struck from −80°C glycerol stocks on to PYE agar and grown at 30°C or room temperature (20-24°C) until colonies appeared after 2-3 days. For static growth experiments, starter cultures (2-10 ml) were inoculated from colonies and grown with aeration overnight at 30°C. Starter cultures were diluted to an optical density at 660 nm (OD_660_) of approximately 0.005 and grown without shaking on the benchtop at room temperature (20-23°C). For experiments requiring repeated sampling through time, I grew cultures with larger surface areas to avoid resampling from the same position. In such experiments, 400 ml of culture was grown in 600 ml Pyrex beakers (9 cm diameter) covered in foil to prevent contamination. In experiments involving only macroscopic inspection of pellicle development, static cultures were inoculated at a similar starting density and grown in test tubes. In preparation of dilute complex medium, the peptone and yeast extract were diluted accordingly from the 1X concentrations of 0.2% and 0.1% w/v, respectively. The MgSO_4_ and CaCl_2_ concentrations were held at constant at 1 and 0.5 mM, respectively.

**Table 1:** Strains used in this work.

| Strain # | Genotype | Source/Reference |
| --- | --- | --- |
| FC19 | *C. crescentus* CB15 | (20, 96) |
| FC20 | *C. crescentus* NA1000 | (46) |
| FC1974 | CB15 $\Delta hfsJ$ | (38) |
| FC1356 | CB15 $\Delta hfiA$ | (38) |
| FC1266 | CB15 $\Delta flgH$ | (31) |
| FC1265 | CB15 $\Delta pilA$ | (31) |
| FC3013 | CB15 $\Delta cpaH$ | (31) |
| FC3084 | CB15 $\Delta pilA \Delta hfiA$ | (31) |
| FC3083 | CB15 $\Delta cpaH \Delta hfiA$ | (31) |
| FC767 | CB15 $\Delta hfsA$ | (46) |
| FC764 | NA1000 *hfsA*<sup>+</sup> | (46) |
| FC766 | NA1000 $\Delta MGE$ | (46) |
| FC3366 | NA1000 $\Delta MGE$ *hfsA*<sup>+</sup> | This work |

### Strain construction

Most of the strains used in this study have been previously constructed and reported (see Table 1). To restore the frameshifted *hfsA* allele on the chromosome of the NA1000 ΔMGE strain I used a standard two-step recombination approach. The pNPTS138-based allele-replacement plasmid carrying the *hfsA*^+^ allele was previously reported, along with methods for using this plasmid (46).

### Sampling from the surface

To capture minimally disturbed cells from the air-liquid interface, I placed the large end of a 1 ml pipet tip on the surface of the static culture. Lifting the tip removed the corresponding segment of the surface layer as a plug (see Figure 1d). I placed the end of the tip carrying the plug sample on a glass slide. I gently applied air pressure to the opposite small end of the tip as I lifted the tip from the slide to ensure complete sample transfer.

### Sampling from the sub-surface

Pipet tips that pass through the pellicle film become coated with film and transfer these cells to anything that directly touches the outside of the tip. To minimize capture of the pellicle when sampling the sub-surface, I submerged a tip 3-4 cm below the liquid surface, expelled air through the tip to blow out any liquid that may have entered the tip and then aspirated several hundred microliters of culture into the tip. After removal of the tip, the culture was rapidly expelled into a sterile tube without making contact between the tip and the tube. If the culture was slowly expelled, the liquid would creep up the side of the tip and capture cells from the pellicle film. Several microliters of the sub-surface sample were placed on a glass slide and covered with a coverslip for microscopic imaging.

### Microscopy

Surface layer plugs placed on glass slides were covered with glass coverslips (see Figure 1e) and imaged using phase contrast with a HCX PL APO 63×/1.4na Ph3 oil objective on a Leica DM5000 upright microscope. Images were captured with a Hamamatsu Orca-ER digital camera using Leica Application Suite X software.

### Fluorescent staining of holdfast

For staining of the holdfast *in situ*, cultures were supplemented with fluorescent 1 ug/ml Wheat Germ Agglutinin conjugated to Alexa Fluor™ 594 (Thermo Fisher) (fWGA) at the time of inoculation. These static cultures were grown under a cardboard box to minimize photobleaching. Samples were collected as above and imaged in phase contrast and fluorescence imaging modes using Chroma filter set 41043.

### Crystal violet staining of pellicle plugs

Surface plugs were placed on glass slides and allowed to stand for 2-4 minutes. After rinsing slides under flowing tap water, the slide was covered with a 0.01% crystal violet solution in water (approximately 1-2 ml to cover the slide). After 3-5 minutes of incubation, the slide was rinsed again and allowed to dry. Stained plugs were photographed with a Nikon 35mm digital camera.

### Oxygen profiling

Oxygen concentrations were measured with a Unisense Field MultiMeter 7614 equipped with a motor controlled micromanipulator and a Clark-type oxygen microelectrode (OX-25; 20-30 μm probe diameter; Unisense). Two point calibrations were performed with air-saturated diH_2_O ([O2] ≈ 283 μM) and a solution of 0.1 M sodium hydroxide, 0.1 M sodium ascorbate (anoxic standard). Calibrations were checked throughout the experiments. Oxygen measurements were performed in 100 μm steps downward starting at the top of the culture. The sensor limit of detection is 0.3 μM O_2_. Profiles for two static cultures for each strain are presented. Measurements were made at the Marine Biological Laboratory (Woods Hole, MA) with equipment loaned to the Microbial Diversity course.

## Acknowledgements

I thank members of the Crosson lab for helpful discussions and feedback. Sean Crosson additionally provided valuable editorial feedback. The dissolved oxygen measurements were performed at the Marine Biological Laboratory (Woods Hole, MA) with equipment loaned to the Microbial Diversity course from Unisense. I thank Bingran Chen for her assistance in making these measurements. This work was supported in part by a grant from the National Institutes of Health (R01GM087353).

