## Supplemental Figures and Movie Legends for "The role of *Caulobacter* cell surface structures in colonization of the air-liquid interface"

Figure S1: Comparison of cells in surface layer pellicle and in the medium sub-surface

Figure S2: Linear arrays of rosettes in pellicle fragments.

### Legends for Supplemental movies

Supplemental Movie 1: Movement of rosette chains under flow

Supplemental Movie 2: *C. crescentus* cells gliding under flow along air-liquid interface I

Supplemental Movie 3: *C. crescentus* cells gliding under flow along air-liquid interface II

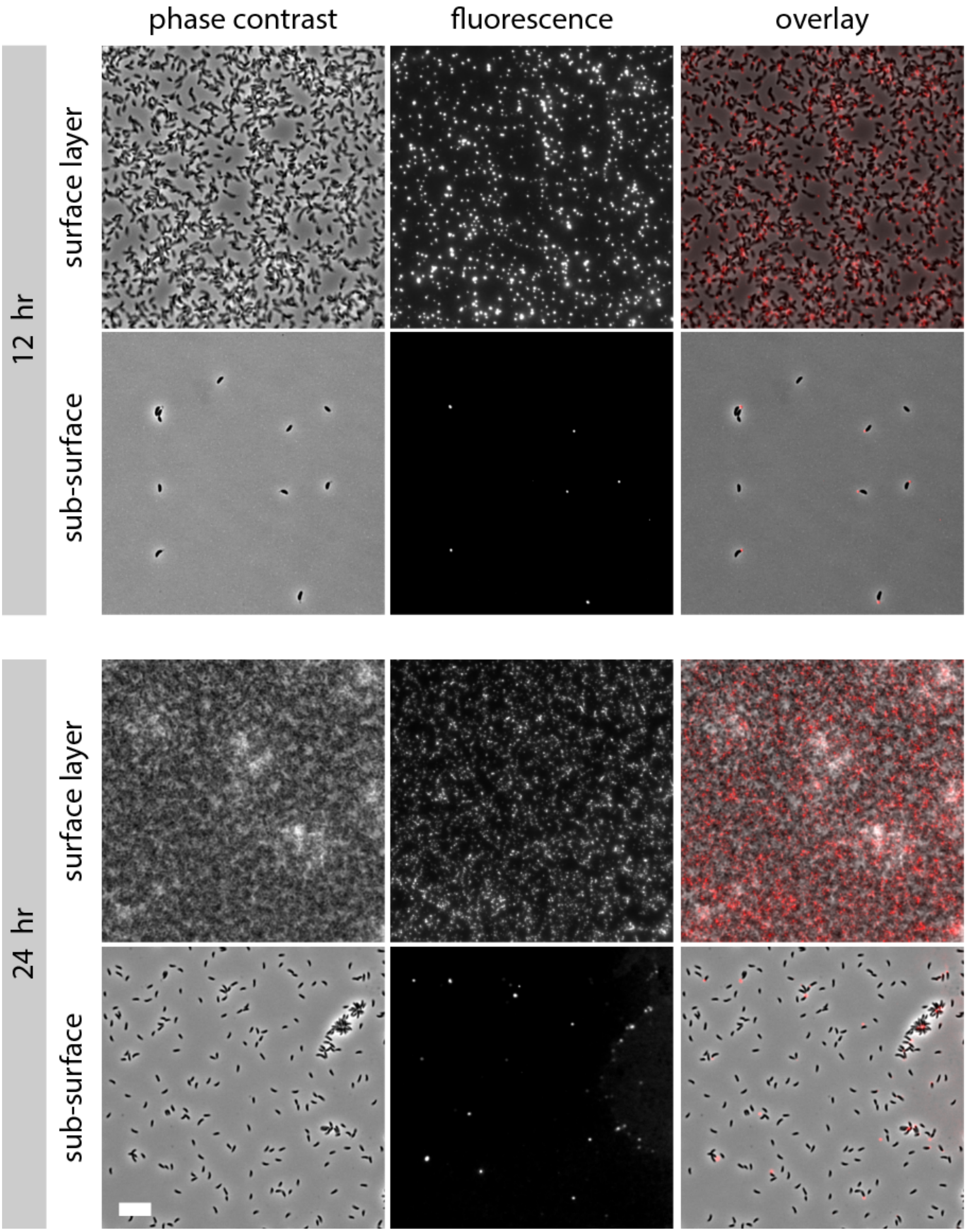


**Figure S1: Comparison of cells in surface layer pellicle and in the medium sub-surface**

Surface film and sub-surface samples of a statically grown CB15 culture supplemented with 1 μg/ml fluorescent wheat germ agglutinin (fWGA) to stain the holdfast. Samples were imaged using phase contrast and fluorescence imaging modes. Scale bar is 10 μm.

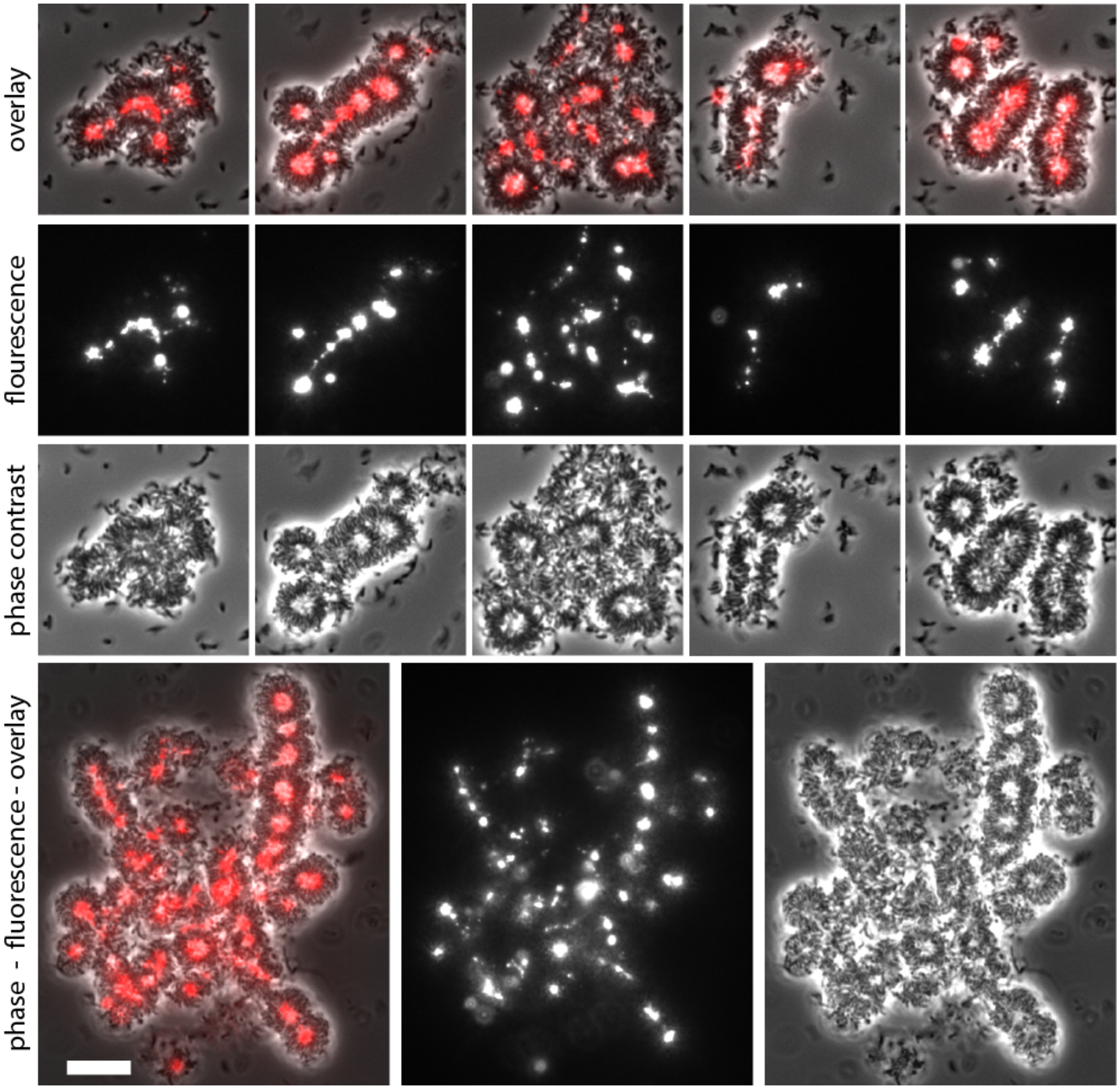


**Figure S2: Linear arrays of rosettes in pellicle fragments.**
Fragments disrupted from the excised pellicle plug shown in the main text Figure 4 were imaged as in Figure 4. Scale bar is 10 μm.

### Legends for Supplemental movies

**Supplemental Movie 1: Movement of rosette chains under flow**
Pellicle plug collected 32 hours after inoculation. The film is in the process of being disrupted by fluid flow on the microscope slide. Individual rosettes and clusters of rosettes are pushed through chains of partially anchored rosettes. Chains of rosettes can be seen moving coordinately under the flow. One example is circled in pink. This phase contrast movie reflects 19 seconds total time (counter in lower left). The edge of an air-bubble crosses the lower left corner. Scale bar is 10 μm.

**Supplemental Movie 2: *C. crescentus* cells gliding under flow along air-liquid interface I**
Phase contrast micrographs of cells from the surface film harvested 32 hours after inoculation near the boundary of a stationary bubble. The liquid flows left to right along the air-liquid interface (indicated by the arrow head). A cluster of cells with long axes oriented perpendicular to the air-liquid interface (phase bright line) glides along this boundary in the direction of flow (highlighted by a pink oval). This cluster resembles a flexible raft that bends with the bumps along the interface. At the end (around 10.5 seconds), a small portion of the cluster gets stuck and breaks away. Individual cells not attached to the air interface tumble across the field of view, pushed by the flow. Movie represents ~ 13 seconds total time (counter in lower left). Scale bar 10 μm.

**Supplemental Movie 3: *C. crescentus* cells gliding under flow along air-liquid interface II**
In this movie, captured from the same sample as supplemental movie 2, many rafts of cells glide around the interface at the edge of an air bubble (right). Flow pushes the cells downward (indicated by arrow head). Moving the sample stage up and down allows visualization of different focal planes along the bubble interface, and also affects the pressure of flow throughout the time sequence. Long axes of cells at the interface are oriented perpendicular to the boundary. Individual cells not attached to the air interface tumble across the field of view, pushed by the flow. A thick mat of cells from the pellicle film occupies the lower left corner. Movie represents nearly 2 minutes total time (counter in lower left). Scale bar 25 μm.
